# Strategies for cellular deconvolution in human brain RNA sequencing data

**DOI:** 10.1101/2020.01.19.910976

**Authors:** Olukayode A. Sosina, Matthew N Tran, Kristen R Maynard, Ran Tao, Margaret A. Taub, Keri Martinowich, Stephen A. Semick, Bryan C. Quach, Daniel R. Weinberger, Thomas M. Hyde, Dana B. Hancock, Joel E. Kleinman, Jeffrey T Leek, Andrew E Jaffe

## Abstract

Statistical deconvolution strategies have emerged over the past decade to estimate the proportion of various cell populations in homogenate tissue sources like brain using gene expression data. Here we show that several existing deconvolution algorithms which estimate the RNA composition of homogenate tissue, relates to the amount of RNA attributable to each cell type, and not the cellular composition relating to the underlying fraction of cells. Incorporating “cell size” parameters into RNA-based deconvolution algorithms can successfully recover cellular fractions in homogenate brain RNA-seq data. We lastly show that using both cell sizes and cell type-specific gene expression profiles from brain regions other than the target/user-provided bulk tissue RNA-seq dataset consistently results in biased cell fractions. We report several independently constructed cell size estimates as a community resource and extend the MuSiC framework to accommodate these cell size estimates (https://github.com/xuranw/MuSiC/).

## Introduction

Homogenate tissues like brain and blood contain a mixture of cell types which can each have unique genomic profiles, and these mixtures of cell types, termed “cellular composition”, can vary across samples^1^. The importance of considering cellular composition within heterogeneous tissue sources has been highlighted in epigenetics research over the past several years^1–3^, as, generally, failure to account for cellular composition when analyzing heterogeneous tissue sources can increase both false positives and negatives^4^. Previous work has identified widespread epigenetic differences between neurons and glia using DNA methylation (DNAm) data^3,5^, and false positives may arise when there are cellular composition differences associated with dissection variability, disease, normal development or any other outcome of interest. For example, loss of neurons (or glia) because of disease may cause spurious loci associations with illness that stem solely from differing cellular compositions between disease states, or cell-type specific biological differences may exist that become more difficult to detect in the presence of unaffected cell types.

Statistical algorithms estimate the relative or absolute amounts of each cell type in the homogenate tissue data. These so called “cellular deconvolution” algorithms have been especially popular using DNAm data^6^ as DNAm levels are constrained between 0 and 1 and are binary within single cells (i.e. individual CpGs are either methylated or unmethylated). These deconvolution algorithms can be classified into two general types, termed “referenced-based” and “reference-free”^6,7^. Reference-free approaches only require as input an estimate of the number of potential cell types in a particular dataset (which can be non-trivial), and return latent components that preferentially capture cellular heterogeneity that can be adjusted for in differential methylation analysis^1,6,8^. However, these approaches do not return fractions of cells and may capture potential batch effects in addition to cellular composition. Conversely, reference-based approaches require cell type-specific genomic profiles for each cell type of interest as an input and return the relative fraction of each input cell type for each queried bulk sample^2^, akin to an *in silico* cell counter. This class of algorithms therefore requires the generation of potentially many pure cell populations, which are typically generated from flow cytometry for applications to DNAm data from bulk tissue.

While DNAm data can generate accurate absolute cell fractions in homogenate brain tissue^3,5,9^, there are several important considerations limiting more widespread application. First, RNA and gene expression profiling has been much more popular in postmortem brain studies, with more samples profiled with RNA sequencing (RNA-seq) than DNAm microarrays or sequencing. Secondly, the two cell classes typically used by DNAm-deconvolution algorithms are likely too broad to identify more subtle differences in dissection variability and potential stereological differences^10,11^. While recent work has extended the number of cell populations that can be isolated by antibodies to separate neurons into their excitatory and inhibitory subclasses and oligodendrocytes from other glia^12^, there are likely very few additional cell types that are possible to isolate using nuclear antibodies for DNAm samples. Researchers have therefore turned to using cell type-specific RNA microarray and sequencing datasets to adapt these reference-based deconvolution algorithms to homogenate RNA-seq samples^7,13–23^. The majority of these studies have focused on tissues other than the brain, which can be freshly obtained and dissociated into individual cells for single cell RNA-seq (scRNA-seq) or be sorted into specific cell populations using flow cytometry for cell type-specific expression profiling. For example, the popular CIBERSORT approach^13^ was designed for blood gene expression microarray data, but has been adapted to RNA-seq datasets in other tissues. Several of the above algorithms have been designed, adapted or implemented for brain tissue, including linear regression followed by quadratic programming using the Houseman algorithm^2,22,23^, non-negative least squares^21^, the support vector machine-based CIBERSORT^24,25^, the empirical Bayes method MIND^18^, and MuSiC, which combines a recursive tree based approach with weighted non-negative least squares for cell type proportion estimation^14^.

However, few of these approaches have validated that the resulting composition estimates are accurate, i.e, are absolutely similar to the true underlying composition, particularly in brain tissue. No approach to our knowledge has quantified the consequences of parameter and algorithm choices when only non-ideal reference data is available (e.g., mismatched tissue type, species, sequencing protocol, etc.), which occurs in almost all applications. Many reference datasets have been constructed from purified cell type-specific RNA-seq data from mouse^26^, or RNA-seq data from sorted or dissociated nuclei in humans^27–31^, and not whole cells, which are typically profiled in homogenate sequencing studies. Gene expression levels are also quantitative within individual cells (and not binary like in DNAm data) and the necessity of absolute expression levels for absolute composition quantification has largely been overlooked.

Here we directly evaluated the absolute accuracy of several popular RNA-seq-based deconvolution strategies using several different reference datasets including a bulk/homogenate dataset with paired DNAm and RNA-seq data from the nucleus accumbens (NAc) from 200+ deceased individuals^32^. We used the DNAm data to estimate absolute neuronal fractions for each sample, and evaluated absolute RNA-based deconvolution accuracy across a variety of scenarios. We first evaluated the effects of using deep single cell RNA-seq (scRNA-seq) from healthy fresh human tissue obtained from surgically resected temporal cortex^33^. This dataset likely produces the most comparable RNA-seq profiles to frozen bulk postmortem tissue, since whole cells were profiled, and 90% of RNA is cytosolic in the cortex^34^. However, this dataset was derived from cells in a cortical brain region. We next produced snRNA-seq data from postmortem human NAc to use as a reference dataset, which results in potentially less comparable nuclear reference profiles but comes from a more comparable brain region. We lastly used cyclic-ouroboros single-molecule fluorescence in situ hybridization (osmFISH) imaging data from the somatosensory cortex region in mouse^35^ to derive important parameters in popular deconvolution algorithms. Together, our results demonstrate that many algorithms are not accurate, even when estimating only two cell classes (neurons and glia), and we offer several strategies to assess and improve accuracy that can be applied across multiple datasets and cell types.

## Results

We motivate this work with a large human postmortem brain genomic dataset from the NAc, a brain region containing functionally distinct cell types critical in reward-processing and addiction^36,37^. Genomic data from this region has been underrepresented in postmortem human brain sequencing studies, which have primarily focused on the frontal cortex^21,38,39^ but its underrepresentation allows us to more comprehensively evaluate the accuracy of cellular deconvolution using potentially imperfect and/or mismatched reference datasets (described below). We dissected homogenate NAc tissue from the ventral striatum (anterior to the optic chiasm) across 223 adult donors and concurrently extracted DNA and RNA from the exact same tissue aliquot (see Methods), which allows for directly comparable cellular composition in each fraction. We profiled genome-wide DNAm with the Illumina Infinium MethylationEPIC microarray and performed reference-based deconvolution to estimate the fraction of neurons in each sample (see Methods). We have previously demonstrated the absolute accuracy of the Houseman deconvolution algorithm^2^ in postmortem human brain DNAm data^5,9^; here we found very high correlation (ρ = −0.949, Figure S1) between the neuronal fraction and the first principal component (PC) of the entire DNAm profile (32.3% of variance explained), which we have shown to be an accurate surrogate of composition in frontal cortex^40^ and blood^1^. The corresponding RNA was sequenced using the Illumina sequencing with RiboZero Gold library preparations (see Methods). This “gold standard” dataset, therefore, has DNAm-derived neuronal composition values and RNA-seq data from 223 samples to explore the accuracy and concordance of many popular cellular deconvolution algorithms.

### Mismatched reference datasets bias deconvolution

We first assessed the accuracy and concordance of four reference-based deconvolution algorithms: Houseman, CIBERSORT, NNLS/MIND, and MuSiC for two cell populations - neurons and non-neurons/glia - in our NAc RNA-seq dataset using recommended default settings (see Methods). We initially used single-cell RNA-seq (scRNA-seq) data from the temporal cortex of eight adult donors obtained during surgical resection generated and described in Darmanis et al., 2015^33^ as the cell type-specific reference profiles for these algorithms. Importantly, these reference data were generated from fresh tissue, which preserved the integrity of the cells and corresponding cytosolic RNA, the predominant fraction of total RNA from brain^34^ profiled in homogenate tissue. Furthermore, these reference profiles provide coverage of entire transcripts (as opposed to only the 3’ ends) using Fluidigm C1 sequencing. Therefore, these expression profiles should be more comparable to bulk brain sequencing studies, with the caveat that the reference dataset was obtained from a different brain region (temporal cortex versus NAc and from living subjects as opposed to postmortem subjects).

We used measures of root mean square error (RMSE) to assess accuracy and squared Pearson correlation coefficients (R^2^) to assess concordance for each algorithm’s estimated neuronal fraction compared to the DNAm-based neuronal fractions (Figure 1). RMSE quantifies the degree of bias, i.e., how much our cell type estimates (RNA composition estimates) deviate from the absolute cell type fractions, with smaller values corresponding to the cellular composition and RNA composition being more similar. R^2^ quantifies the amount of information our estimates contain about how the absolute cell type fractions vary in the population being studied, i.e., how much variability of the cell type fractions, across individuals, is captured by our composition estimates. Houseman (Figure 1A), MuSiC (Figure 1B), and NNLS produced concordant (high correlation; Houseman R^2^ = 0.51, p < 2.20E-16; MuSiC R^2^ = 0.56, p < 2.20E-16; NNLS R^2^ = 0.54, p < 2.20E-16) but biased (high RMSE, >0.35) neuronal fraction estimates. CIBERSORT produced more discordant (moderate correlation; R^2^ = 0.25, p = 5.13E-03) neuronal fraction estimates (Figure 1C), but with less bias (low RSME, 0.09). We found that CIBERSORT, compared to either MuSiC or the Houseman RNA approach, was the most accurate. However, its estimates provided the least information (R^2^ value) about the variability of the estimates based on DNAm data. In comparing the R^2^ metric across the three approaches, we found that MuSiC provided the most information about the observed variability of the observed cell type proportions among the 223 individuals but was the most biased. These results suggest that all four of these approaches overestimated the proportion of neurons in bulk brain tissue, even under the simplest application to deconvoluting two distinct cell populations. However, it was unclear how much algorithm parameters and reference dataset differences (in regards to technology and brain region) contributed to the performance of these methods.

**Figure 1:**
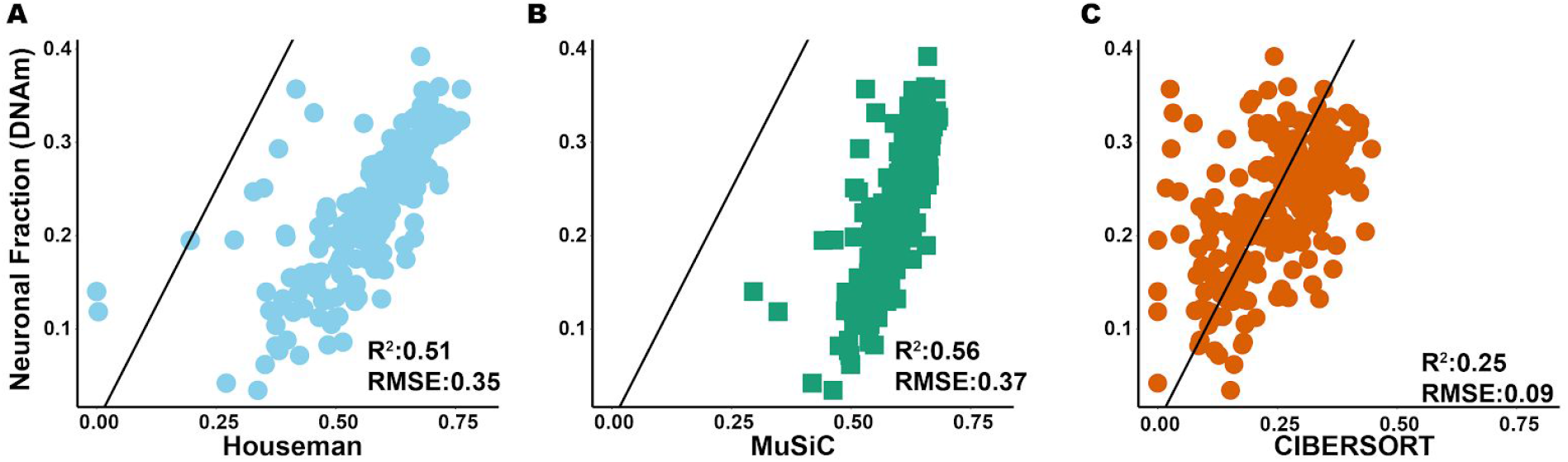
Deconvolution in bulk NAc data using gene expression profiles from the temporal cortex (Darmanis); Scatter plots showing the estimated neuronal proportions across the 223 individuals using the Houseman approach for DNAm reference vs neuronal proportions estimated using (**A**) the Houseman approach with scRNA reference, (**B**) MuSiC with default settings and scRNA reference data, and (**C**) CIBERSORT with scRNA reference data.

### Methods for reducing bias in cellular deconvolution

Many of the above deconvolution strategies have several parameters whose adjustment could reduce the observed bias (i.e. maximize accuracy) and increase the concordance between these neuronal fractions. The MuSiC algorithm particularly has an interpretable “cell size” (see Methods) parameter used in the deconvolution process. Different cell types could have more or less absolute RNA abundance, for example if they were larger or smaller, or if they were more or less transcriptionally active. We hypothesized that the overestimation of neuronal fractions resulted from neurons being larger and more transcriptionally active. However, this “cell size” parameter, regularly defined by the algorithm as the average expression level for a given cell type summed across genes, is estimated directly from the reference cell type-specific RNA-seq profiles by default (see Methods). However, some scRNA-seq (or snRNA-seq) library preparation and sequencing strategies, like the Fluidigm C1 system, may normalize cDNA libraries to the same concentration prior to sequencing, which will remove potential variability in RNA abundances across cell types. We therefore sought to use external data to better estimate these cell size parameters (Table 1) and assessed the resulting effects on cellular deconvolution accuracy.

**Table 1:** Cell sizes used for deconvolution.

| Cell type | NAc 50<br>genes (UMIs) | NAc 25<br>genes (UMIs) | NAc all<br>genes (UMIs) | Temporal<br>cortex,<br>Darmanis<br>et al.<br>(Counts) | osmFISH cell<br>Area ($\mu\text{m}^2$) | osmFISH<br>nRNA<br>(intensity) |
| --- | --- | --- | --- | --- | --- | --- |
| Glial | 710.63 | 453.24 | 5763.55 | 12879.73 | 90.87 | 180.46 |
| Neuronal | 4513.58 | 2793.54 | 29884.65 | 18924.66 | 122.96 | 198.86 |
| Neuronal<br>/glial<br>ratio | 6.35 | 6.16 | 5.19 | 1.47 | 1.35 | 1.1 |

UMI, unique molecular identifier, counts= $\log_2(\text{cpm}+0.5)$ , intensity = sum of probe intensities across 24,048 genes.

First, we used external ouroboros single-molecule fluorescence in situ hybridization (osmFISH) data from mouse somatosensory cortex^35^ to construct two different types of cell size parameters for the MuSiC algorithm (as data from NAc did not exist). We extracted the estimates of both cell size (via their provided segmentations) and total RNA abundance (via the sum of all gene fluorescence signal) aggregated across neuronal and non-neuronal cell types. We subsequently utilized these estimates as proxies for cell size in human RNA-seq data when deconvoluting neuronal fractions. In these data, comparing neurons to non-neurons, neurons were both larger (123 vs 91 μm^2^, p < 2.20E-16) and had more total RNA (199 vs 180 intensity, p = 1.73E-05) as we observed in the estimated cell size in the MuSiC algorithm using the Darmanis dataset (18,925 vs 12,880 normalized counts). We did not observe any improvement in the concordance (osmFISH cell area R^2^ = 0.55, p < 2.2E-16; osmFISH totalRNA R^2^ = 0.54, p < 2.2E-16) or accuracy (osmFISH cell area RMSE = 0.39; osmFISH totalRNA RMSE = 0.43) of the estimated cell type fractions when we compared our results from default settings to those based on applying cell size proxies using mouse data (Figure S2 A-B). These results may not be particularly surprising, given the numerous differences between mouse and human morphology, and the different brain regions profiled.

We then generated snRNA-seq dataset from 2 postmortem NAc donors and 4,169 total nuclei to produce more comparable cell type specific cell size (see Methods) parameters and reference expression profiles (see Methods). First, we used the NAc reference dataset at the single nucleus level and ran the MuSiC algorithm with default settings, which used both NAc-based cell sizes and expression profiles, to deconvolute neuronal fractions (Figure 2A). We confirmed that, on average, neurons had more total RNA than non-neurons using this NAc snRNA-seq dataset (103 vs. 72 unique molecular identifiers [UMIs] per gene, p < 2.2E-16). Furthermore, while there was a high correlation among neuron-specific gene expression effects across the NAc and temporal cortex (Darmanis et al.) reference profile datasets, we observed genes with different magnitudes of effects based on differential expression results between neuronal and non-neuronal cell types (Figure S3). When using both the NAc-based cell size and gene expression reference profiles, we observed a substantial improvement in both the concordance and the RMSE for the estimated neuronal fractions compared to using the temporal cortex dataset only. However, the estimates were still biased, and this bias increased as the neuronal fraction across individuals increased, suggesting that the NAc-based cell sizes together with the estimated abundance may be incorrectly characterizing the true underlying neuronal expression level for these individuals. Eliminating the cell size parameter resulted in similarly reduced concordancies in both the temporal cortex and the NAc reference datasets, but increased accuracy only using the NAc reference dataset (Table 2). This implies that the underlying broad cellular composition was well captured by the gene “abundance” information for a matched brain region.

**Figure 2:**
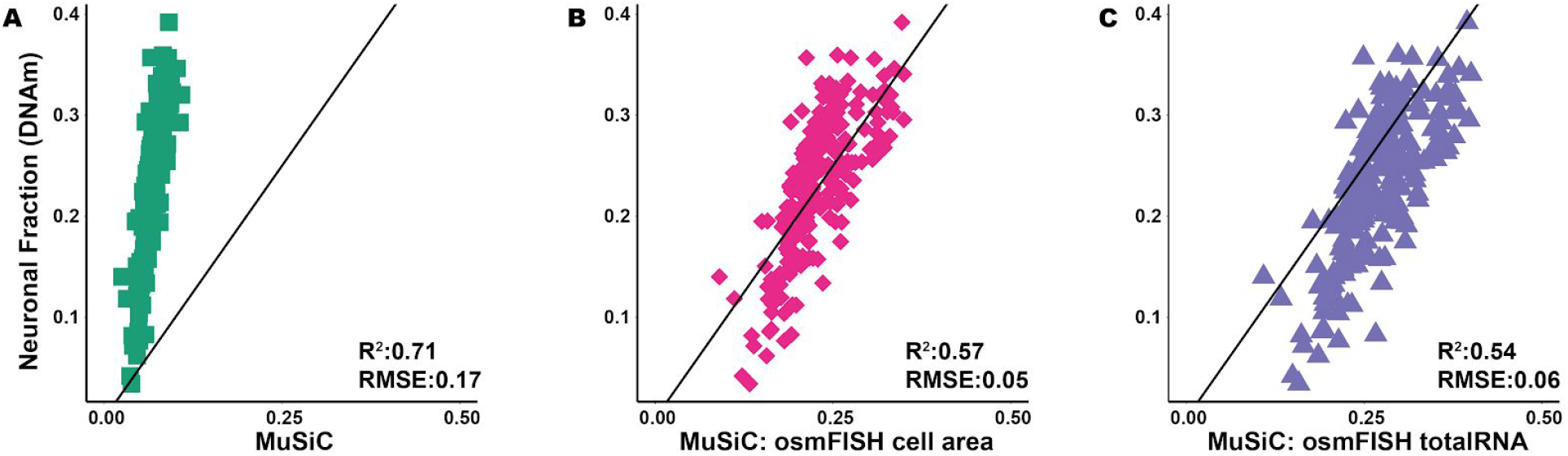
Deconvolution in bulk NAc data based on a single nucleus RNAseq (snRNA-seq) reference dataset from the same brain region. Scatter plots comparing the estimated neuronal proportion obtained for each individual using the Houseman approach with DNAm reference dataset vs neuronal proportions obtained using (**A**) MuSiC with default settings and a snRNA-seq NAc reference dataset, (**B**) MuSiC based on a snRNA-seq NAc reference dataset with cell sizes for each cell type estimated using osmFISH cell area (mouse), and (**C**) MuSiC based on a snRNA-seq NAc reference dataset with cell sizes for each cell type estimated using osmFISH total RNA abundance (mouse) per cell type.

**Table 2:** Bias and concordance results for deconvolution of bulk NAc data using each cell size and gene expression reference dataset.

| Method | Cell size | Reference dataset |  |  |  |
| --- | --- | --- | --- | --- | --- |
|  |  | scRNA-seq in temporal cortex (Darmanis et al.) |  | snRNA-seq in NAc |  |
| | | Concordance ($R^2$) | Accuracy (RMSE) | Concordance ($R^2$) | Accuracy (RMSE) |
| MuSiC | None | 0.54 | 0.45 | 0.54 | 0.08 |
|  | scRNAseq in temporal cortex (Darmanis et al.) | 0.56 | 0.37 | 0.58 | 0.05 |
|  | snRNAseq in NAc | 0.59 | 0.08 | 0.71 | 0.17 |
|  | snRNAseq in NAc (Top 25 genes) | 0.59 | 0.06 | 0.71 | 0.18 |
|  | snRNAseq in NAc (Top 50 genes) | 0.59 | 0.05 | 0.72 | 0.18 |
|  | osmFISH-Cell area | 0.55 | 0.38 | 0.57 | 0.05 |
|  | osmFISH-Total RNA | 0.54 | 0.43 | 0.54 | 0.06 |
| CIBERSORT | N/A | 0.25 | 0.09 | N/A | N/A |
| NNLS | N/A | 0.54 | 0.40 | 0.72 | 0.22 |

We then combined different estimates of cell size parameters (NAc snRNA-seq versus osmFISH) and gene expression reference profiles (NAc snRNA-seq versus temporal cortex scRNA-seq) and assessed the effects on deconvolution accuracy in bulk NAc RNA-seq data. When running MuSiC using the estimates of cell size based on osmFISH data with the NAc expression reference profiles, we observed further improvements in the bias of the estimated cell type fractions but saw a minimal difference in the concordance (Figure 2B and C). Surprisingly, when we used only the Darmanis cell type-specific expression levels, the best (least biased and most concordant) deconvolution results were produced using cell sizes estimated from NAc snRNA-seq data, with improvements in both the concordance and the RMSE (Figure 3A). Specifically, when we compared the R^2^ and the RMSE estimates to those observed under the default setting for the Darmanis reference with the mismatched brain region, we see a small (6% relative change, p = 3.3E-02) increase for the concordance and a substantial (78% relative change, p < 1E-04) decrease for the RMSE. We further refined the NAc cell size estimates using sets of the top 25 and 50 cell type discriminating genes (see Methods), which slightly improved our estimates of the absolute cell type fractions (Figure 3B and 3C). Both the concordance (7% relative change, p = 3.5E-02) and RMSE (86% relative change, p < 1E-04) improved even more when compared to the default approach using a mismatched reference dataset. Across all approaches, the most accurate (least biased) result occurred when we used cell sizes estimated from Darmanis scRNAseq data and gene expression from NAc snRNA-seq data, while the most concordant results were observed when we used NAc snRNAseq data exclusively.

**Figure 3:**
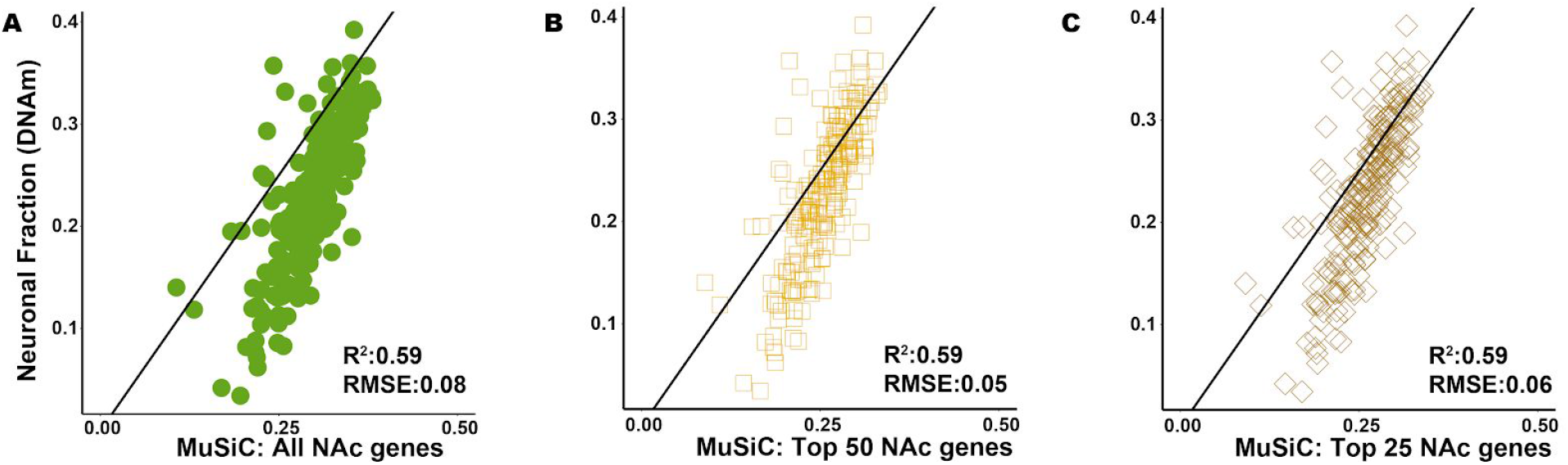
Deconvolution in bulk NAc data using gene expression profiles from the temporal cortex (Darmanis et al) and different estimates of cell size. Scatter plots comparing the neuronal fraction estimated for each individual using DNAm data and the Houseman method vs neuronal fractions based on scRNA-seq data and estimated using MuSiC with (**A**) cell-size estimated using all genes expressed in the NAc snRNAseq reference dataset, (**B**) cell-size estimated using the top 50 cell type discriminating genes in the NAc snRNAseq reference dataset, and (**C**) cell-size estimated using the top 25 cell type discriminating genes in the NAc snRNAseq reference dataset.

In summary, when we used a region-matched appropriate dataset - NAc snRNA data - as the reference, or to derive estimates of the cell size, we observed that estimates of the cell type proportions generally improved (Table 2, Figure 2 A and Figure 3A-C). In settings where we had a mismatched reference dataset (e.g., mismatched on brain region or species), incorporating estimated cell sizes obtained from the matched brain region (NAc) provided the best result in metrics for both concordance and accuracy, and we slightly improved these metrics when we refined the gene sets used to estimate the cell sizes.

## Discussion

Statistical deconvolution strategies have emerged over the past decade to estimate the proportion of various cell populations in homogenate tissue sources like blood and brain from both gene expression and DNAm data. Our results together suggest that many existing RNA deconvolution algorithms estimate the RNA composition of homogenate tissue, e.g. the amount of RNA attributable to each cell type, and not the cellular composition, which relates to the underlying fraction of cells. This was evident by the consistent overestimation of larger and more transcriptionally active neuronal cells. We have identified that incorporating cell size parameters into RNA-based deconvolution algorithms can successfully recover cellular fractions in homogenate brain RNA-seq data. We have lastly shown that using both cell sizes and cell type-specific gene expression profiles from brain regions other than the target/user-provided bulk tissue RNA-seq dataset consistently resulted in overestimating neuronal fractions. We have developed an extension of the MuSiC framework^14^ that allows for the incorporation of independent cell size estimates, and have further provided cell size estimates for human brain (shown in Table 1) as a part of the package: https://github.com/xuranw/MuSiC.

Characterizing cellular heterogeneity is especially important in human brain, where the underlying cell types can have diverse functions and disease associations that could be missed in studies of bulk tissue (Michels et al. 2013). Here we show that RNA-based deconvolution for just two cell populations - neurons and non-neurons - largely fails to estimate the underlying cellular composition of bulk human brain tissue across a variety of algorithms and strategies. We quantified the diverse range of neuronal fractions estimated by several popular algorithms to better understand the effects of reference cell type-specific expression profiles and differences in cell size and/or activity profiles on deconvolution. We specifically examined the common scenario of performing RNA deconvolution using cell type-specific reference datasets that can be fundamentally different from user-provided homogenate tissue target datasets, for example differing in profiled brain region, sequencing technology and/or cellular compartment. These problems are likely magnified in human brain tissue compared to suspended cells like blood, where deconvolution strategies are more easily validated against true cell fractions obtained by routine complete cell counts^13^. We lastly emphasize caution when performing RNA-based deconvolution using many cell types (i.e., more finely-partitioned cell classes) without having the ability to validate cell counts on at least a subset of samples.

We therefore offer several recommendations for performing RNA-based deconvolution in bulk human brain gene expression data, particularly when aiming to identify cellular, and not RNA, composition.

1. Providing estimates of cell size for each reference cell type improves the concordance and reduces bias when performing RNA deconvolution to estimate cellular fractions. Biologically-motivated and valid external estimates of cell-size improve the accuracy of the estimated cell type fractions, even when gene expression profiles for reference cell populations are obtained from other brain regions (Figure 3). The exact biological interpretation of these estimated cell sizes, particularly when estimated across species, is arguably unclear, but likely relates to correcting for absolute RNA abundance and differences in transcriptional activity between cell populations. Regardless of the method used for deriving cell sizes, neurons consistently had more RNA than glia. We note that our recommended strategies for estimating cell size have only been assessed for broad classes of cell types, and further work is needed to validate extensions to more stratified subclasses of cells.
2. The concordance and bias improvements using full-length single cell sequencing from a different brain region (temporal cortex), rather than single nuclei RNA-seq from the target brain region (NAc) highlighted the importance of comparability between reference gene expression profiles and the homogenate tissue expression levels. While previous reports have identified high correlation between nuclear and cytosolic gene expression levels in both bulk^41^ and single cell^30,42^ resolution, comparable absolute (and not relative) expression levels are seemingly important for the accuracy of these RNA-based cellular deconvolution algorithms. There further is an experimental design tradeoff between profiling more nuclei (1000s) using 3’ technologies like 10x Genomics Chromium Single Cell Gene Expression compared to profiling fewer nuclei (or cells, 100s) using full-length sequencing technologies like SMART-seq if researchers wish to generate their own reference profiles.
3. Using reference cell type-specific expression profiles from comparable brain regions as the bulk RNA-seq target dataset is important, and can especially greatly increase the concordance of these RNA deconvolution strategies with neuronal fractions.

The choice of maximizing accuracy (by minimizing bias) versus increasing concordance in assessing these algorithms is an important consideration, particularly when generating custom expression reference profiles is prohibitive (Table 2). These two objectives largely relate to whether the goal of RNA deconvolution is to estimate cell fractions (and maximize accuracy) or RNA fractions (and maximize concordance). Estimation of RNA fractions (by maximizing concordance) may be sufficient to control for potential confounding due to composition differences between outcome groups^4^. We note this can also be accomplished using “reference-free” deconvolution^6^ or through the estimation of potentially sparse principal components^1,8^ that control for relative differences in cellular composition. However, estimation of cellular fractions (and maximizing accuracy) is arguably more useful, both for assessing human brain tissue dissection during data generation and to identify cell type-specific effects when using these cellular fractions in downstream differential expression analyses^43^.

Together, our results demonstrate that many RNA deconvolution algorithms do not produce accurate cellular fractions when estimating only two cell classes (neurons and non-neurons). We offer several strategies and corresponding software to assess and improve accuracy that can be applied across multiple datasets and cell types.

## Methods

### Bulk NAc Data Generation and Processing

Data generation and processing were described extensively in Markunas et al. 2019^32^. Briefly, the nucleus accumbens (NAc) was dissected under visual guidance using a hand-held dental drill. Samples were obtained from the ventral striatum, anterior to the optic chiasm, at the level where the NAcforms a bridge between the putamen and the head of the caudate. DNA and RNA were concurrently extracted from dissected tissue using the Qiagen AllPrep DNA/RNA Mini Kit (Cat No./ID: 80204).

NAc DNA was profiled with the Infinium MethylationEPIC microarray using the manufacturer’s protocol. Raw idat files were processed and normalized using the minfi Bioconductor package^44^ using stratified quantile normalization^45^. Resulting neuronal fractions were estimated using the minfi estimateCellCounts function^1^ using sorted reference data from the DLPFC for neurons and non-neurons^5,46^ using the Houseman algorithm^2^.

NAc RNA was subjected to RNA-seq library preparations using the Illumina RiboZero Gold kits and sequenced using 2×100bp paired end reads on an Illumina HiSeq 3000.

### Reference Datasets

Darmanis (Darmanis et al., 2015)

scRNA-seq data for 58,037 genes and 556 cells were obtained for brain samples across 8 individuals, as described previously (Darmanis et al., 2015). We filtered this dataset by removing cells based on embryonic samples and retaining cells from one of the following five cell types; Neuronal, Oligodendrocyte progenitor cells (OPC), Astrocytes, Oligodendrocytes, and Microglia. We also removed genes that had no expression for all cells in the reference dataset or did not show any expression in the bulk dataset (i.e., mean and variance zero). In total, we used 265 cells for this reference and 24,048 genes to estimate the cell type proportions for the 223 samples with bulk NAc data.

### Single-nucleus RNA-seq data generation and processing in nucleus accumbens

We performed single-nucleus RNA-seq (snRNA-seq) on nucleus accumbens (NAc) tissue from two donors using 10x Genomics Single Cell Gene Expression V3 technology. Nuclei were isolated using a “Frankenstein” nuclei isolation protocol developed by Martelotto et al. for frozen tissues^27,47–50^. Briefly, ~40mg of frozen NAc tissue was homogenized in chilled Nuclei EZ Lysis Buffer (MilliporeSigma) in a glass dounce with ~15 strokes per pestle. Homogenate was filtered using a 70um-strainer mesh and centrifuged at 500xg for 5 minutes at 4°C in a benchtop centrifuge. Nuclei were resuspended in the EZ lysis buffer, centrifuged again, and equilibrated to nuclei wash/resuspension buffer (1x PBS, 1% BSA, 0.2U/uL RNase Inhibitor). Nuclei were washed and centrifuged in this nuclei wash/resuspension buffer three times, before labeling with DAPI (10ug/mL). The sample was then filtered through a 35um-cell strainer and sorted on a BD FACS Aria II Flow Cytometer (Becton Dickinson) at the Johns Hopkins University Sidney Kimmel Comprehensive Cancer Center (SKCCC) Flow Cytometry Core. Gating criteria hierarchically selected for whole, singlet nuclei (by forward/side scatter), then for G_0_/G_1_ nuclei (by DAPI fluorescence). A null sort was additionally performed from the same preparation to ensure nuclei input was free of debris. Approximately 8,500 single nuclei were sorted directly into 25.1uL of reverse transcription reagents from the 10x Genomics Single Cell 3’ Reagents kit (without enzyme). Libraries were prepared according to manufacturer’s instructions (10x Genomics) and sequenced on the Next-seq (Illumina) at the Johns Hopkins University Transcriptomics and Deep Sequencing Core.

We processed the sequencing data with the 10x Genomics’ Cell Ranger pipeline, aligning to the human reference genome GRCh38, with a reconfigured GTF such that intronic alignments were additionally counted given the nuclear context, to generate UMI/feature-barcode matrices. We used R package Seurat^51^ for raw feature-barcode quality control, dimensionality reduction (PCA), choosing the top 30 PCs as the optimal dimensions for clustering. We performed graph-based clustering with the default Louvain approach, taking a computed K-nearest neighbors graph as input, which were then annotated with well-established cell type markers for nuclear type identity^31^. We also used Seurat’s implementation of non-linear dimensionality reduction techniques, t-SNE and UMAP, simply for visualization of the high-dimensional structure in the data, which complemented the clustering results (Supp Figure 4). With the five broad cell type annotations (neurons, oligodendrocytes, oligodendrocyte precursors, astrocytes, and microglia) of nuclear clusters, we identified unbiased cluster-driving genes (with Seurat’s *‘FindAllMarkers* ()’ function, using the Wilcoxon rank-sum test), that were upregulated in each cell type/cluster, compared to all other nuclei. Using the same set of 24,048 genes, we have 4,169 high-quality nuclei in this reference, evenly distributed across donors. The top 50- and top 25-per-cell-type gene sets had 247 and 125 genes, respectively, which included many cell type marker genes used for annotation.

### Estimation procedures

#### HOUSEMAN

This algorithm (Houseman et al., 2012) uses a linearly constrained quadratic optimization approach with additional non-negative constraints on the parameters. The linear constraint does not require that the sum of all coefficients equal one. This allows the possibility of unknown cell types in case the specification is not comprehensive. It was implemented using the minfi R Bioconductor package (Aryee et al. 2014).

#### MuSiC

The MuSiC (Wang et al., 2019) approach models the relationship between the relative abundance of gene g in the bulk RNA-seq data and the mean expression level of the same gene in the reference dataset for a given individual. The relationship is provided below

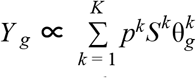

Where k = 1,…,K is the index of the cell types, *p^k^* is the proportion of cells from cell type k, and 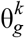 is the relative abundance for the gth gene with respect to the kth cell type. *S^k^* is the cell size parameter and is defined as the average number of total mRNA molecules for cell type k. By default, *S^k^* is estimated automatically by MuSiC. For the deconvolution method comparisons that assessed cell size impact on neuronal cell type proportion estimation, *S^k^* was derived from one of multiple data sources (Table 1) using 1) default settings 2) osmFISH or 3) the average number of total mRNA molecules for cell type k using only the top 25 or 50 most discriminating genes per cell type. We defined “most discriminating” as genes with the smallest p-values and fold change >0.25, relative to other cell types. All estimation was carried out using the MuSiC package in R.

#### CIBERSORT

CIBERSORT uses a machine learning approach called nu-support vector regression (Newman et al., 2015; Schölkopf et al., 2000) and requires at least 2 input datasets to work. The first is a signature matrix that identifies the set of genes that are informative for the deconvolution procedure. The second is a bulk RNA-seq dataset to estimate cell type proportions.

The signature matrix depends on the tissue of interest. We generated a custom signature matrix. Using the Darmanis reference dataset, we generated both a reference sample file (gene-by-cell matrix) and a phenotype classes file (cell type-by-dummy variable identifying the cell type for each cell) and used the default setting (<u>CIBERSORT</u>) to obtain a custom signature gene expression matrix. The specified false discovery rate (FDR) threshold used to include genes in the signature matrix was 0.30 (i.e. q = 0.30, default). Using this signature matrix, we then performed deconvolution on our bulk NAc RNA-seq data. As suggested in the documentation for CIBERSORT (<u>CIBERSORT</u>), we disabled quantile normalization for our RNA-seq data.

#### NNLS/MIND

This is a simple linear regression with non-negativity constraints on the parameter estimates. The estimated fractions are then the value of each parameter estimate divided by the sum of all parameter estimates across cell types. MIND (https://github.com/randel/MIND) uses NNLS to estimate cell type fractions.

## Acknowledgements

The authors thank the authors of MuSiC; X. Wang, J. Park, K. Susztak, N.R. Zhang, and M. Li for their technical assistance and valuable insights. K.M., S.A.S., D.B.H., and A.E.J. were partially supported by R01DA042090

## Supplementary Information

**Supplementary Table 1:** Proportion of cell types in datasets used to generate cell sizes or as reference data

| Cell type | NAC | Darmanis | osmFISH |
| --- | --- | --- | --- |
| Non-neuronal | 0.83 | 0.51 | 0.31 |
| Neuronal | 0.17 | 0.49 | 0.69 |

**Supplementary Figure 1.**
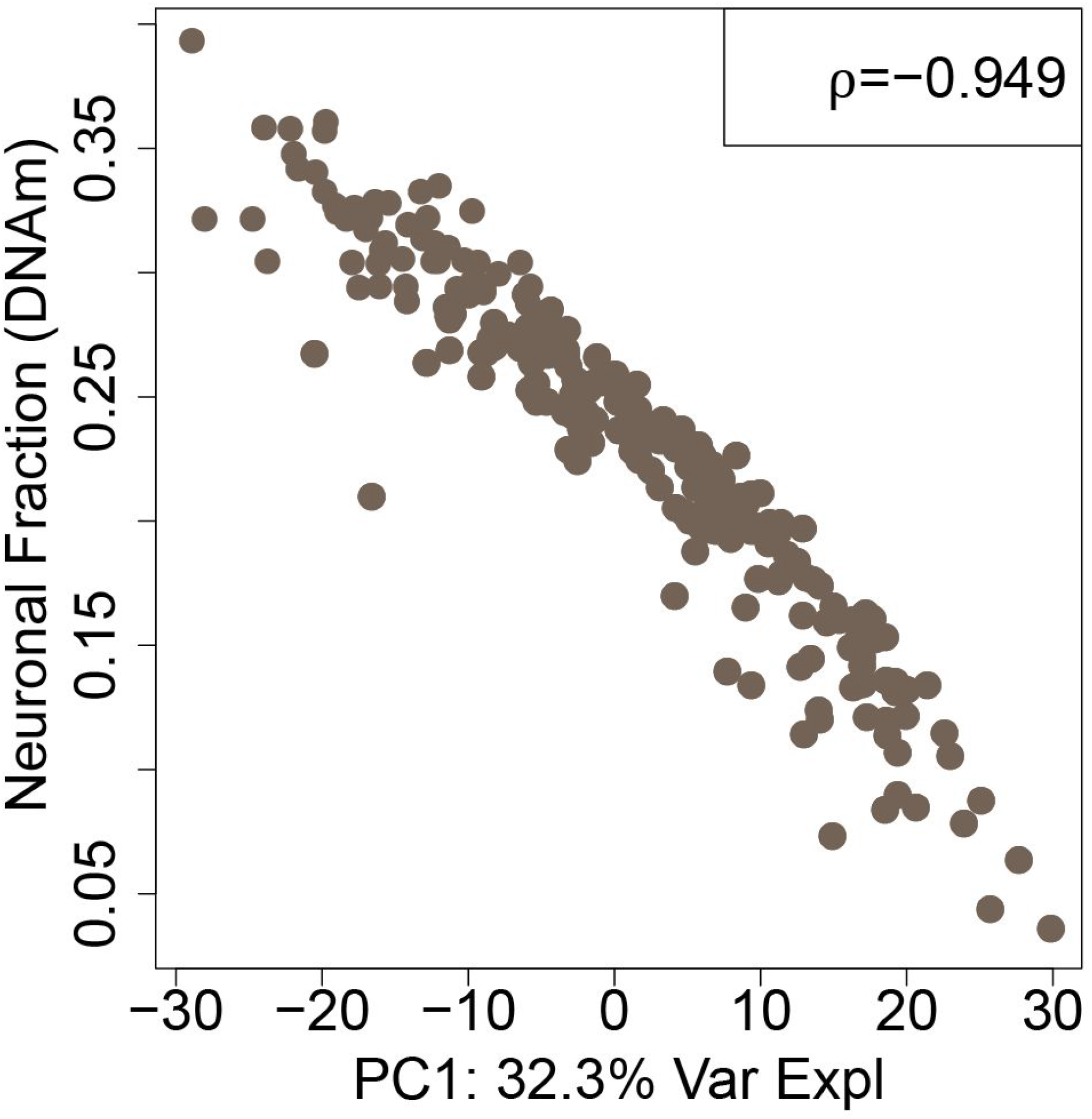
(DNAm estimated neuronal fractions vs PC1): Scatter plot of neuronal fractions estimated using the Houseman approach with a DNAm reference vs the first principal component estimated from the bulk RNA-Seq data.

**Supplementary Figure 2:**
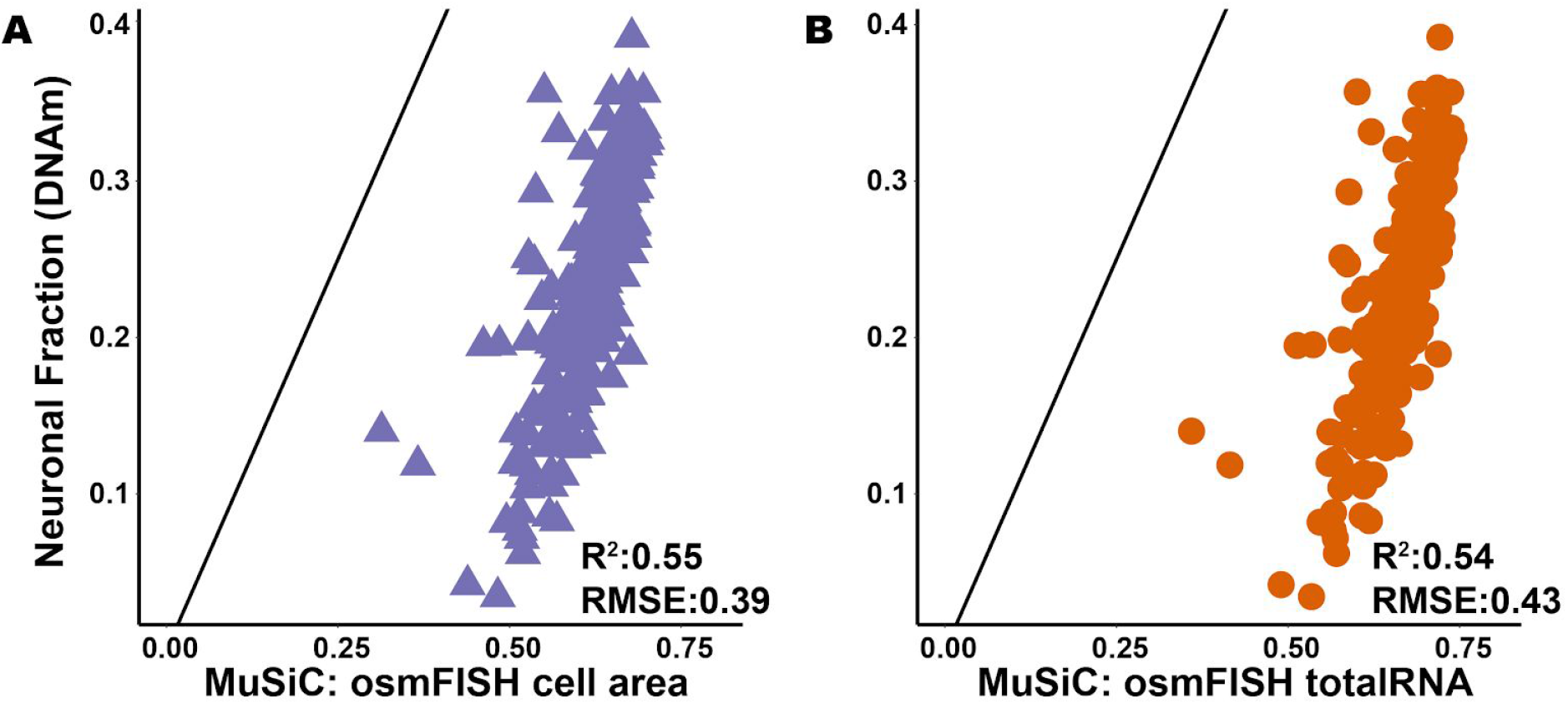
Deconvolution in bulk NAc data using gene expression profiles from the temporal cortex (Darmanis et al) with cell size estimates derived using mouse samples (osmFISH estimates of cell size). Scatter plots comparing neuronal fraction estimated for each individual using DNAm data and the Houseman method vs neuronal fractions based on scRNA-seq data and estimated using MuSiC with (**A**) osmFISH cell area as cell size, and (**B**) osmFISH total RNA molecule count as cell size

**Supplementary Figure 3:**
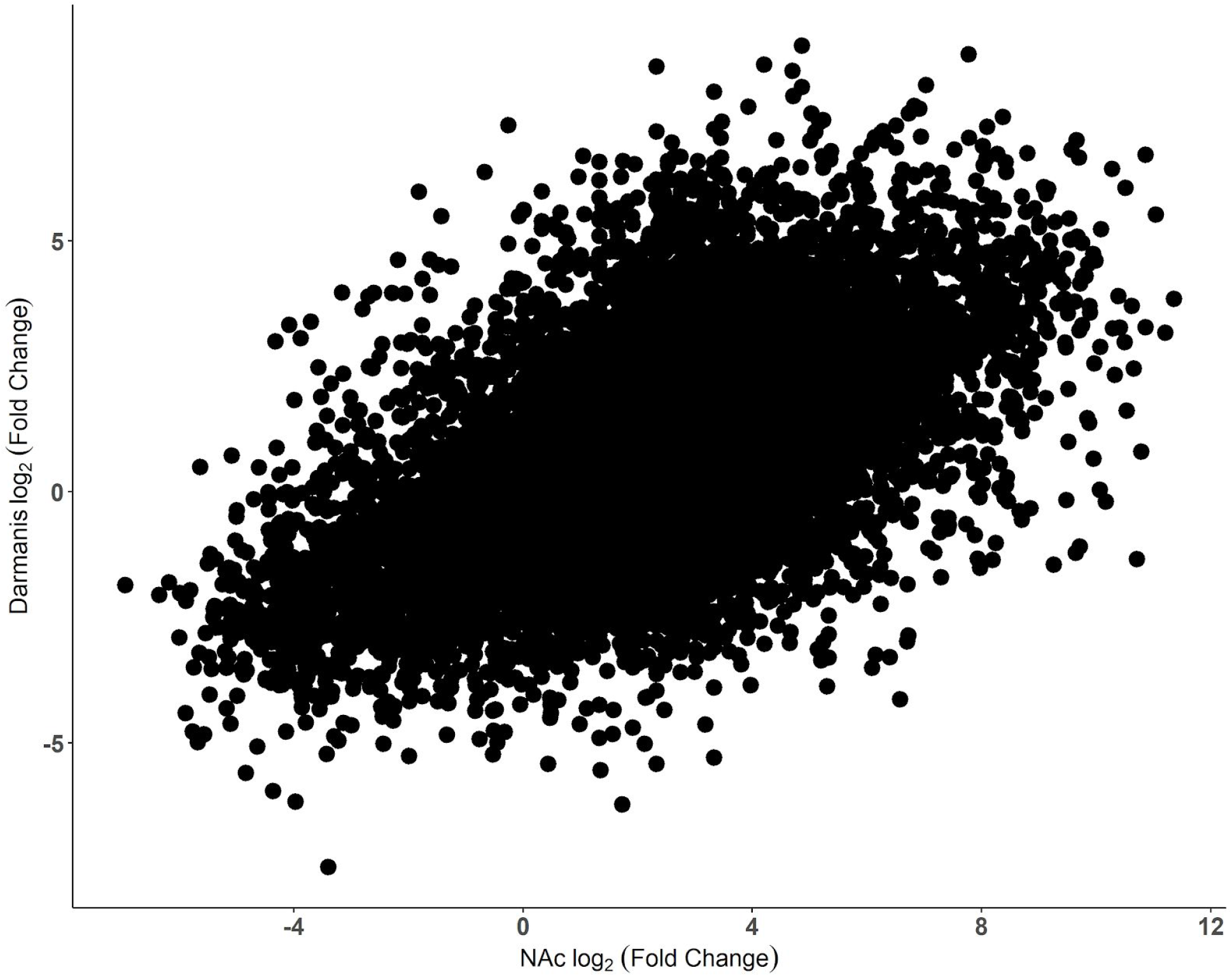
Neuronal enrichment of gene expression in scRNA-seq from temporal cortex and snRNA-seq from nucleus accumbens. Scatter plot shows the relationship, based on *log*_2_ (fold change) comparing neuronal to glial, between the Darmanis reference dataset (y-axis) and the NAc reference dataset (x-axis). Each dot represents an estimated *log*_2_ (fold change) for a given gene.

**Supplementary Figure 4:**
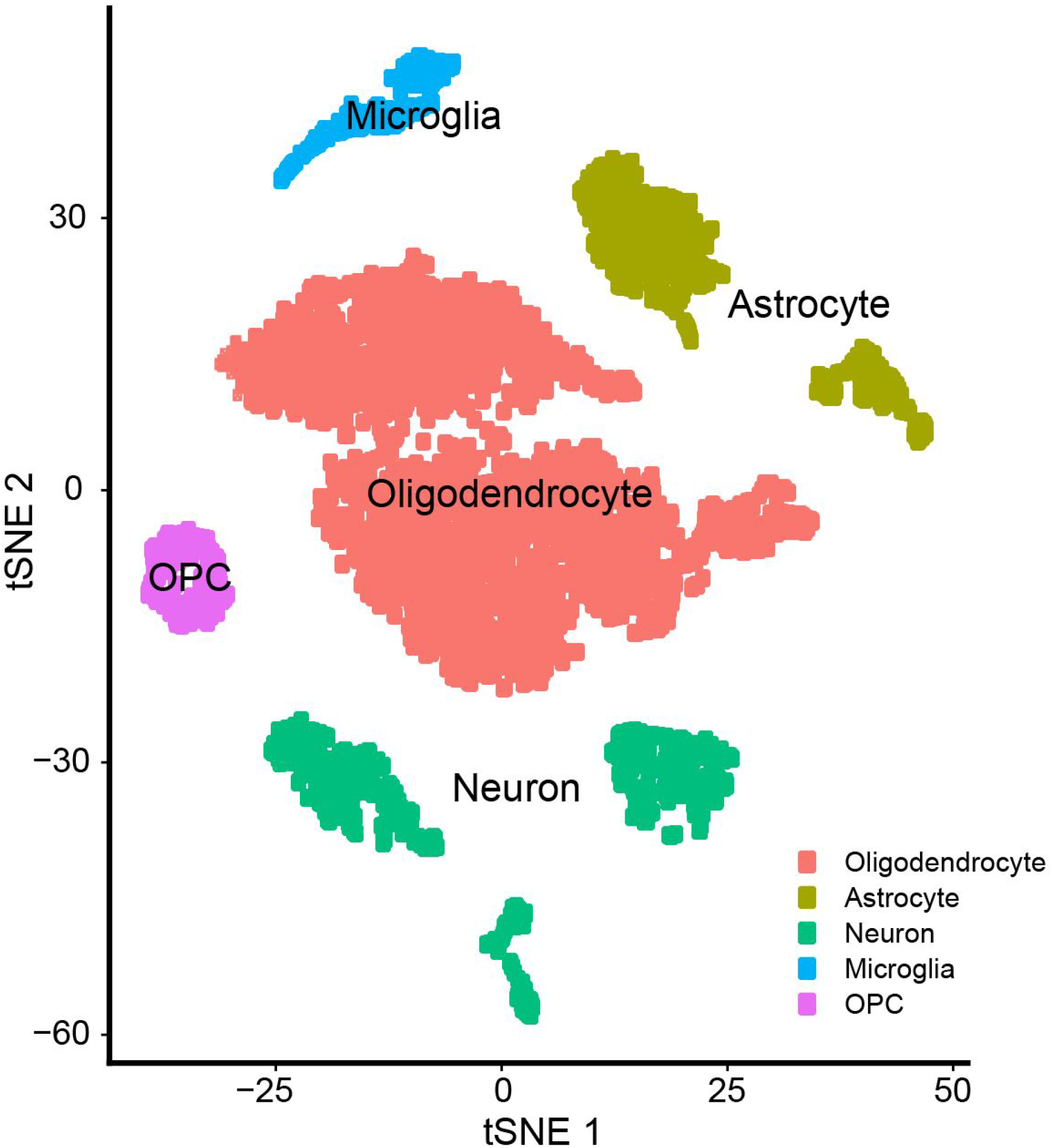
t-distributed stochastic neighbor embedding (t-SNE) of single-nucleus RNA-seq data from the two postmortem NAc samples, representing the 4,169 high-quality nuclei after processing. Nuclei are colored by cell type annotation after graph-based clustering, which shown here is largely in agreement with t-SNE coordinates. OPC represents Oligodendrocyte progenitor cell

